# Mid-pregnancy placental transcriptome in a model of placental insufficiency with and without novel intervention

**DOI:** 10.1101/2024.06.05.597621

**Authors:** Rebecca L. Wilson, Baylea N. Davenport, Helen N. Jones

## Abstract

Fetal growth restriction (FGR) affects between 5-10% of all live births. Placental insufficiency is a leading cause of FGR, resulting in reduced nutrient and oxygen delivery to the fetus. Currently, there are no effective in utero treatment options for FGR, or placental insufficiency. We have developed a gene therapy to deliver, via a non-viral nanoparticle, *human insulin-like 1 growth factor* (*hIGF1*) to the placenta as potential treatment of placenta insufficiency and FGR. Using a guinea pig maternal nutrient restriction (MNR) model of FGR, we aimed to understand the transcriptional changes within the placenta associated with placental insufficiency that occur prior to/at initiation of FGR, and the impact of short-term *hIGF1* nanoparticle treatment. Using RNAsequencing, we analyzed protein coding genes of three experimental groups: Control and MNR dams receiving a sham treatment, and MNR dams receiving *hIGF1* nanoparticle treatment. Pathway enrichment analysis comparing differentially expressed genelists in sham-treated MNR placentas to Control revealed upregulation of pathways associated with degradation and repair of genetic information and downregulation of pathways associated with transmembrane transport. When compared to sham-treated MNR placentas, MNR + *hIGF1* placentas demonstrated changes to genelists associated with transmembrane transporter activity including ion, vitamin and solute carrier transport. Overall, this study identifies the key signaling and metabolic changes occurring in the placenta contributing to placental insufficiency prior to/at initiation of FGR, and increases our understanding of the pathways that our nanoparticle-mediated gene therapy intervention regulates.

**Statements and Declarations:** *Competing Interests:* Authors declare no conflicts of interest.

## Introduction

Fetal growth restriction (FGR) is the second leading cause of perinatal mortality, and affects between 5-10% of all live births [1]. In developing countries more than 20% of babies are born FGR, but there are currently no treatment options [2]. Individuals with FGR are at greater risk of cardiovascular and renal problems, metabolic problems such as type II diabetes, and other disorders in adulthood because of their suboptimal in-utero environment and developmental programming [3]. The principal cause of FGR is placental insufficiency, leading to reduced nutrients and oxygen delivery to the fetus [4]. To address this, our lab has developed a potential treatment for placental insufficiency and in-turn FGR using a placenta targeted nanoparticle gene therapy to deliver *human insulin-like 1 growth factor* (*hIGF1*). We have now shown in multiple animal models of FGR that our treatment is able to increase nutrient transporters, alter vascular factors, and treat reduced fetal growth [5-9]

Placental insufficiency is defined by reduced capability of the placenta to transfer nutrients and oxygen to the fetus either due to maldevelopment or malfunction. This leads to a hypoxic environment and downregulation of metabolic demands to protect the fetus as much as possible. This reduced nutrient transfer, metabolism and oxygen, however, leads to the decreased growth of the fetus[10]. Within the placenta, the environmental changes associated with placenta insufficiency include alterations to the villous (human) or labyrinth (guinea pig) vasculature, decreased trophoblast proliferation and differentiation, reduced nutrient transfer capabilities and decreases in both growth factors and their related signaling pathways that lead to these physiologic changes [11]. In human transcriptome studies comparing Control and FGR placentas, analysis identified differential expression of genes related to responses to reactive oxygen species, ions, increased protein translation, and receptor kinase signaling [12]. While these changes were reported at the end of term, placental insufficiency begins long before this and early diagnosis is crucial for both the mother and fetus. However, there are no biomarkers to detect FGR in early pregnancy, and diagnosis based on fetal growth trajectory occurs in the second half of pregnancy [13]. Because of this it is imperative to have animal models to understand the mechanisms of placental insufficiency during pregnancy.

The guinea pig model offers many advantages and similarities to humans. Compared to common rodent models, the guinea pig’s gestation is longer (∼65 days), and compared to humans both the placenta and fetus reach similar developmental milestones through gestation and after birth [14]. Guinea Pigs also have a haemomonochorial placenta and deep trophoblast invasion like that of humans [15, 16]. To model FGR in these guinea pigs we employ the maternal nutrient restriction (MNR) model. This is a well-established model that uses a reduced percentage of normal food intake to create placenta insufficiency through undernutrition [17]. The MNR model has been shown many times to create a similar placental environment to human FGR cases and alters similar signaling cascades, such as the IGF axis, ultimately leading to lowered fetal weight [18, 19]. Using this model, we can understand the impact of FGR on the placenta at various timepoints to understand the implications in early and mid-pregnancy to further our understanding of underlying mechanisms.

Insulin-like Growth Factor 1 (IGF1) is actively produced by the placenta throughout the entirety of pregnancy, but in cases of FGR, IGF1 levels have been shown to be decreased [20, 21]. Trophoblasts synthesize and secrete IGF1 to regulate nutrient transport, angiogenesis, and trophoblast invasion and proliferation [21]. Because of its importance in placenta function and development throughout the entirety of pregnancy and its reduction in FGR, we have developed a gene therapy to deliver *hIGF1* to the placenta as potential treatment of placental insufficiency and FGR. To deliver the *hIGF1* gene under the control of a trophoblast-specific promoter, we use a self-forming nanoparticle and ultrasound guided intraplacental injection [5, 6]. We have previously published that we are able to increase fetal capillary volume density and reduce the interhaemal distance between maternal and fetal circulation in the MNR placenta with *hIGF1* nanoparticle treatment, showing our treatment’s potential to positively impact placental capability for nutrient and oxygen transport [22].

With placental insufficiency leading to hypoxic and deteriorative environments, cells within this environment respond to these stressors. These cell stress responses can create vast changes in regulation of gene expression and determine cell fate. With this study we aimed to identify transcriptional changes in the placenta associated with MNR and placental insufficiency, as well as the alterations made by *hIGF1* nanoparticle treatment. Utilization of this knowledge can then be used to better understand mechanisms underlying placental insufficiency at a cellular signaling level and our ability to correct these changes, thus, paving the way for more effective *in utero* treatment option for FGR in the future.

## Methods

### Nanoparticle Formation

Nanoparticles were formed by complexing plasmids containing *hIGF1* under the control of trophoblast-specific promoter *CYP19A1* with a non-viral PHPMA_115_-b-PDMEAMA_115_ co-polymer as described previously [22]. Maternal and fetal safety have been demonstrated previously in numerous animal models in [6, 22-24].

### Animals

Experiments were approved by Cincinnati Children’s Hospital Medical Center (Protocol 2017-0065). Female Dunkin Hartley guinea pigs (Charles River) were housed in a controlled environment (22°C / 50% humidity / 12hr light-dark cycle) and provided food (Labdiet 5025: 27% protein, 13.5% fat and 60% carbohydrate as % of energy) and water *ad libitum*. After a 2-week acclimation period, females were weighed and randomly assigned to Control *ad libitum* fed diet or maternal nutrient restriction (MNR) diet described in [19, 25]. Timed mating, pregnancy confirmation, and ultrasound-guided intraplacental nanoparticle injections were performed as previously described in [22]. At gestational day 30-33, females underwent *hIGF1* nanoparticle or PBS sham injection. 5 days later animals were sacrificed (GD35-38). Major maternal and fetal organs were collected, fetal sex was determined and placentas were collected. Sub-placenta/decidua was separated from placenta labyrinth and tissues were flash frozen in liquid nitrogen and stored at -80°.

### RNAsequencing Library Preparation and Sequencing

RNA was isolated from frozen placentas (Control n = 8; MNR n = 8; MNR + *hIGF1* n = 8) using the RNeasy mini kit (Qiagen), including DNAse treatment, following standard protocols. RNA (RIN>5) was sent to the University of Florida’s Institute for Interdisciplinary Center for Biotechnology Research (ICBR) where they generated libraries from 1.5 µg RNA using the Illumina Stranded mRNA Prep Kit following manufacturers specifications. RNA sequencing was performed on the NovaSeq 6000. There were ∼50,000,000 reads per sample and libraries were clustered for paired-end sequencing. Raw and processed sequencing data is available on NCBI Geo (GSE269097).

### RNAsequencing Differential Analysis

Short reads were trimmed using trimmomatic (v 0.36) [26]. Quality control on the original and trimmed reads was performed using FastQC (v 0.11.4) and MultiQC [27]. Reads were aligned to the *Cavia porcellus* transcriptome using STAR (v 2.7.9a), and transcript abundance was quantified using RSEM (v 1.3.1) [28, 29]. Differential expression (DE) analysis was performed using DESeq2. Genes with a Log2 fold change of 1.0 in both directions, and a raw p-value of 0.05 difference between Control vs MNR, Control vs MNR + *hIGF1*, and MNR vs MNR + *hIGF1* were selected for pathway analysis.

### Pathway Analysis

Heat map was generated using the Broad Institute’s Morpheus software. Gene lists were sorted by p-value prior to software upload. For those genes within multiple groups, p-values were ordered according to that of the “Control vs MNR” group’s p-value.

Pathway enrichment analysis was performed using ToppFun (ToppGene Suite V31 [30]). Differentially expressed genes were separated into upregulated genes (positive log changes) and downregulated genes (negative log changes) and analyzed separately. P values were calculated using the Hypergeometric Probability Mass Function and false discovery rate corrected using Benjamini–Hochberg methods.

## Results

Initially, potential confounding of fetal sex between placentas from males and females was assessed. Principal component analysis (PCA) on normalized expression data showed no separation of samples by fetal sex and therefore, data generated from male and female placentas were combined for further analysis (Figure 1A).

**Fig. 1.**
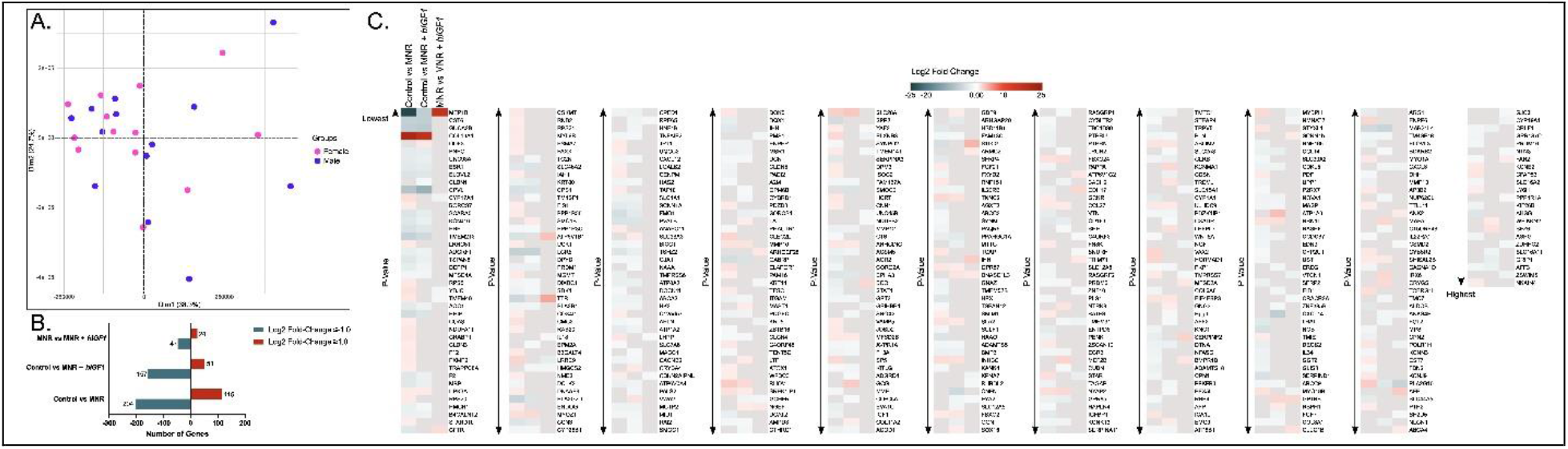
Differential gene expression analysis in the guinea pig placenta at mid-pregnancy. **A**. Principal component analysis (PCA) on normalized expression data showed no separation of samples by fetal sex. n = 12 female and 12 male. **B**. Number of upregulated and down regulated differentially expressed genes comparing Control and maternal nutrient restriction (MNR), Control and MNR + *hIGF1* nanoparticle treatment, and MNR and MNR + *hIGF1*. n = 8 Control, 8 MNR and 8 MNR + *hIGF1*. **C**. Heat map of differentially expressed genes across Control, MNR and MNR + *hIGF1*.

### Differentially expressed genes

Between Control and sham-treated MNR placentas, 319 differentially expressed protein-coding genes were identified (Figure 1B, 1C). 115 of which were upregulated while 204 genes were downregulated. The most upregulated genes in MNR placentas included *Col16A1* (log2 FC: 21.42) an integral collagen extracellular matrix component, and *Cyp17A1* (log2 FC: 1.79) a cytochrome p450 gene important for cellular metabolism (Figure 1C). The most downregulated genes in MNR placentas included: *Mep1B* (log2 FC: -24.06) an extracellular protease involved in connective tissue homeostasis; *Cpvl* (log2 FC: -8.04) a carboxypeptidase involved in post-translational modifications; *Guca2B* (log2 FC: -6.76) involved in salt and water homeostasis, and *Cst6* (log2 FC: -6.46) a cystatin protease inhibitor.

In MNR + *hIGF1* placentas, the number of differentially expressed genes compared to Controls were 208, 51 of which were upregulated and 157 were downregulated. Within this group, the genes with the largest fold change were: *Col16A1* (log2 FC: +17.60) and *Atp1A3* (log2 FC: +3.41) a P-type cation transport ATPases, and *Cpvl* (log2 FC: -11.14), *Guca2B* (log2 FC: -7.43), and *Mep1B* (log2 FC: -6.85).

Comparing MNR + *hIGF1* placentas to sham-treated MNR placentas resulted in identification of 71 differentially expressed genes. *Mep1B* (log2 FC: 17.20) and *Ttr* (log2 FC: 4.80) a carrier protein predicted to be involved in glucose homeostasis were the most upregulated, whilst *Pla2G10* (log2 FC: -6.07) a phospholipase A2 family member with roles in the production of inflammatory lipid mediators, *Lrrc9* (log2 FC: -2.60) a paralogue of *Lrguk* which is involved in kinase activity and *Star* (log2 FC: -2.35) a regulator of steroid hormone synthesis were downregulated. The full list of differentially expressed genes with p-values and fold changes are included in the supplementary materials.

### Maternal nutrient restriction results in reduced representation of pathways involved in nutrient transport and increased enrichment of DNA repair pathways

Pathway enrichment analysis of differentially expressed genes in sham-treated MNR placentas compared to Control revealed changes associated with degradation and repair of genetic information and cellular metabolism (Table 1). Differentially expressed genelists increased in MNR placentas were enriched for pathways including enzyme inhibitor activity (FDR: 1.37E-01, p-value: 3.48E-03), phosphatase inhibitor activity (FDR: 6.06E-02, p-value: 2.50E-04), regulation of apoptotic DNA fragmentation (FDR: 5.27E-01, p-value: 4.22E-03) and DNA catabolic processes (FDR: 5.27E-01, p-value: 5.20E-03). Genelists of down-regulated genes in MNR placentas were enriched for pathways including transport of small molecules (FDR: 5.29E-03, p-value: 3.74E-05), ion channel transport (FDR: 6.03E-03, p-value: 5.98E-05), regulation of IGF1, IGF1 transport, and uptake by IGF binding proteins (FDR: 1.85E-03, p-value: 2.64E-06).

**Table 1.**
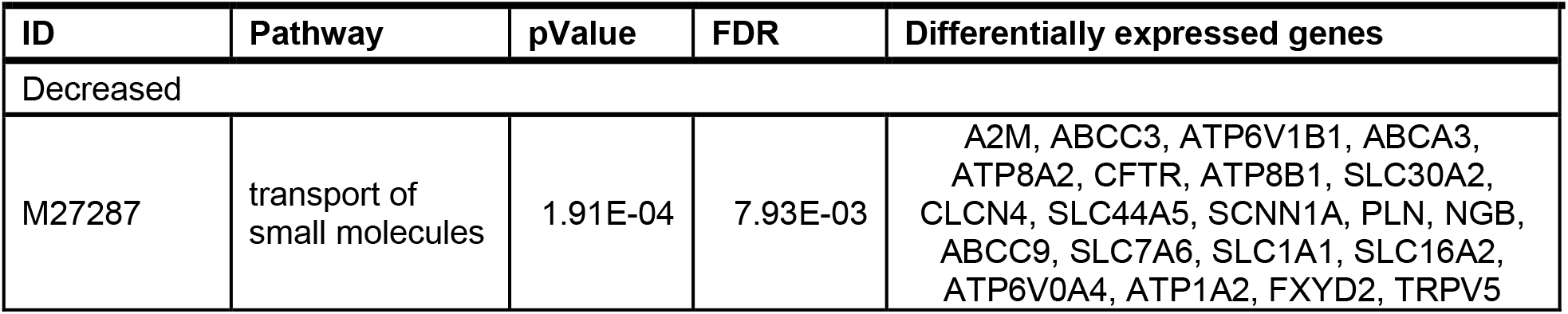
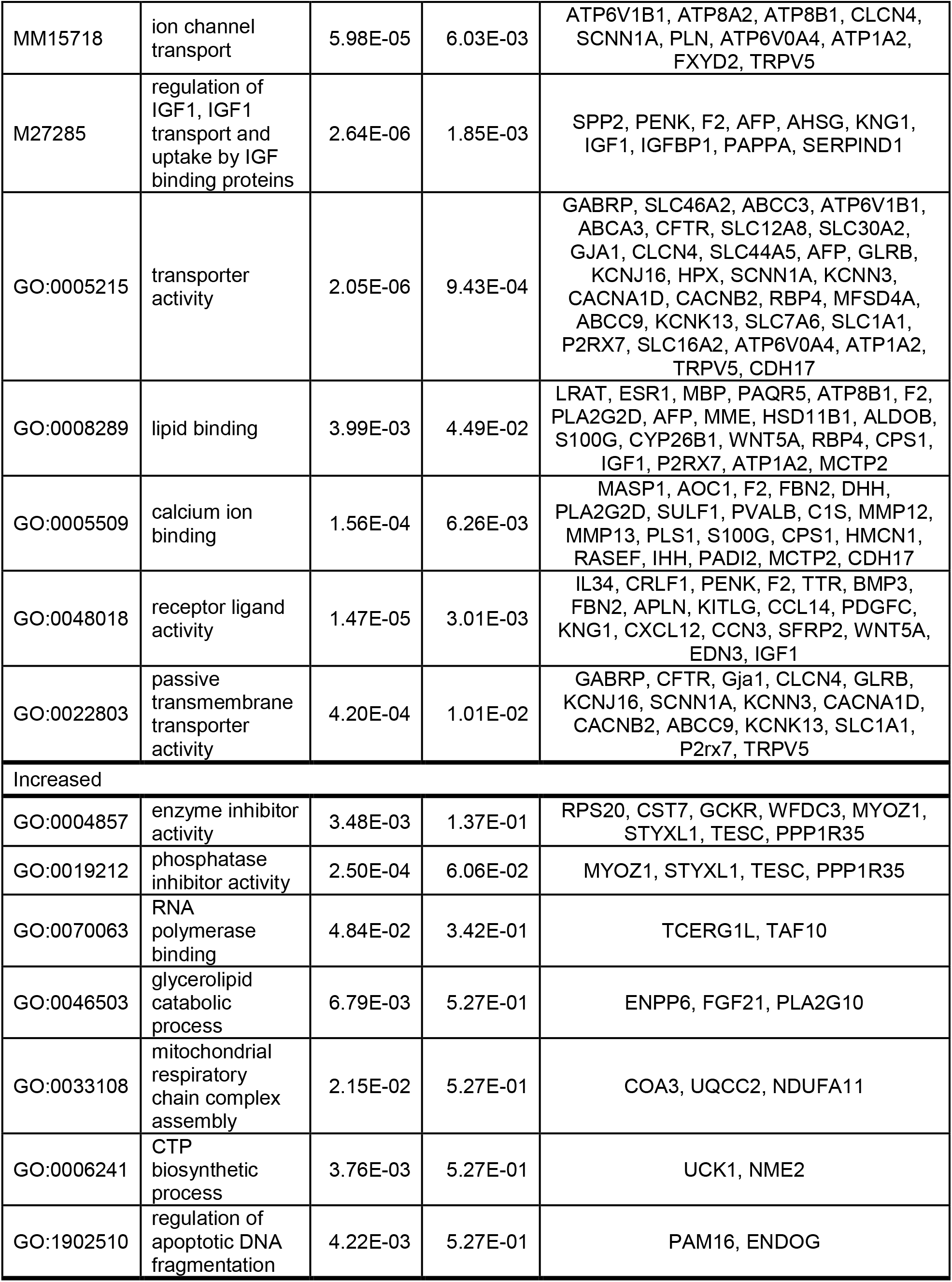
Pathway enrichment analysis of differentially expressed genes in sham treated maternal nutrient restricted (MNR) placentas compared to Control.

Similarly, differentially expressed genelists that were decreased in MNR + *hIGF1* placentas compared to Control were enriched for pathways relating to transporter activities (transporter activity: FDR: 8.11E-03, p-value: 1.22E-04; channel activity: FDR: 8.11E-03, p-value: 8.61E-05) and growth factor activity (FDR: 9.67E-04, p-value: 1.45E-06) (Table 2). However, unlike sham-treated MNR placentas, differentially expressed genelists that were increased in MNR + *hIGF1* placentas compared to Control were enriched for pathways relating to positive regulation of phosphorylation (FDR: 2.82E-01, p-value: 2.46E-02) and positive regulation of kinase activity (FDR: 2.47E-01, p-value: 8.75E-03)(Table 2).

**Table 2.** Pathway enrichment analysis of differentially expressed genes in *IGF1* nanoparticle treated maternal nutrient restricted (MNR + *IGF1*) placentas compared to Control.

| ID | Pathway | pValue | FDR | Differentially expressed genes |
| --- | --- | --- | --- | --- |
| Decreased |  |  |  |  |
| GO:0005215 | transporter activity | 1.22E-04 | 8.11E-03 | GABRP, SLC46A2, CACNB2, ABCA3, CYBRD1, TMC7, SLC2A8, NUP62CL, ATP8A2, MFSD4A, ABCC9, KCNK13, ATP8B1, PEX5L, SLC1A1, SLC12A5, GJA1, CLCN4, GLRB, ATP6V0A4, KCNJ16, SCNN1A, SLC16A11, ATP1A2 |
| GO:0048018 | receptor ligand activity | 2.98E-05 | 4.90E-03 | MACC1, EREG, CRLF1, CXCL14, F2, OGN, BMP3, KITLG, APLN, FGF7, CCL14, PDGFC, CXCL12, CCN3, CXCL8 |
| GO:0015267 | channel activity | 8.61E-05 | 8.11E-03 | GABRP, CACNB2, TMC7, NUP62CL, ABCC9, KCNK13, PEX5L, SLC1A1, SLC12A5, GJA1, CLCN4, GLRB, KCNJ16, SCNN1A |
| GO:0022839 | monoatomic ion gated channel activity | 4.04E-04 | 1.58E-02 | GABRP, CACNB2, TMC7, ABCC9, KCNK13, PEX5L, CLCN4, GLRB, KCNJ16, SCNN1A |
| GO:0140326 | ATPase-coupled intramembrane lipid transporter activity | 1.36E-03 | 3.35E-02 | ABCA3, ATP8A2, ATP8B1 |
| MM14563 | metabolism | 1.60E-02 | 2.01E-01 | DPYD, CACNB2, AMPD3, FMO1, AOC1, HK2, CYP26B1, GCHFR, HMGCS2, PFKFB3, CPS1, DCN, OGN, ELOVL2, PNPO, PPP1R3C, PLA2R1, PLA2G2D, HPGD, B4GALNT2, ACSM5, PLA2G10, HAS2, GPT2 |
| M27287 | transport of small molecules | 3.49E-03 | 8.87E-02 | CYBRD1, SLC2A8, ATP8A2, ABCC9, ATP8B1, CUBN, SLC1A1, SLC12A5, CLCN4, ATP6V0A4, SCNN1A, ATP1A2, PLN |
| Increased |  |  |  |  |
| GO:0042327 | positive regulation of phosphorylation | 2.46E-02 | 2.82E-01 | LAT, INHBC, NTRK3, GCG, LTF, GDF9 |
| GO:0033674 | positive regulation of kinase activity | 8.75E-03 | 2.47E-01 | LAT, NTRK3, GCG, LTF, GDF9 |
| GO:0019722 | calcium-mediated signaling | 3.84E-03 | 2.14E-01 | LAT, MYOZ1, SLA2, TREML1 |
| GO:0071902 | positive regulation of protein serine/threonine kinase activity | 2.19E-02 | 2.70E-01 | NTRK3, LTF, GDF9 |

*Short term hIGF1 nanoparticle treatment results in increased representation of pathways involved in transmembrane transporter activity and reduced enrichment of cellular metabolism pathways hIGF1* nanoparticle treatment of the MNR placenta resulted in enrichment of pathways such as transmembrane transporter activity including ion transport, vitamin transport and SLC-mediated transport when compared to sham treated MNR placentas (Table 3). Differentially expressed genelists that were increased in MNR + *hIGF1* placentas were enriched for pathways including active transmembrane transporter activity (FDR: 3.82E-03, p-value: 5.66E-05), inorganic cation transmembrane transporter activity (FDR: 3.02E-02, p-value: 5.53E-03), vitamin D binding (FDR: 1.35E-02, p-value: 9.78E-04), and ABC-type transporter activity (FDR: 1.66E-02, p-value: 1.65E-03). Differentially expressed genelists that were decreased in MNR + *hIGF1* placentas compared to sham-treated MNR were enriched for pathways including lipid biosynthetic process (FDR: 1.54E-01, p-value: 8.40E-03), and hormone metabolic process (FDR: 1.26E-01, p-value: 4.03E-03).

**Table 3.** Pathway enrichment analysis of differentially expressed genes in *IGF1* nanoparticle treated maternal nutrient restricted (MNR + *IGF1*) placentas compared to sham MNR.

| ID | Pathway | pValue | FDR | Differentially expressed genes |
| --- | --- | --- | --- | --- |
| Decreased |  |  |  |  |
| GO:0008610 | lipid biosynthetic process | 8.40E-03 | 1.54E-01 | PLA2G10, HMGCS2, RPE65, STAR, CYP17A1, CYP19A1 |
| GO:0042445 | hormone metabolic process | 4.03E-03 | 1.26E-01 | STAR, CYP17A1, CYP19A1 |
| GO:0097192 | extrinsic apoptotic signaling pathway in absence of ligand | 1.42E-02 | 1.88E-01 | NGF, EYA2 |
| GO:0008211 | glucocorticoid metabolic process | 1.62E-03 | 1.20E-01 | STAR, CYP17A1 |
| Increased |  |  |  |  |
| GO:0022857 | transmembrane transporter activity | 5.66E-05 | 3.82E-03 | SCNN1B, ATP6V1B1, MFSD2A, ATP6V1G2, ABCA4, AFP, CDH17, CFTR |
| GO:0022890 | inorganic cation transmembrane transporter activity | 5.53E-03 | 3.02E-02 | SCNN1B, ATP6V1B1, MFSD2A, ATP6V1G2 |
| GO:0019842 | vitamin binding | 9.78E-04 | 1.35E-02 | ABCA4, S100G, AFP |
| GO:0140359 | ABC-type transporter activity | 1.65E-03 | 1.66E-02 | ABCA4, CFTR |
| GO:0000902 | cell morphogenesis | 6.66E-04 | 3.95E-02 | CLEC1B, MFSD2A, BMPR1B, ARHGEF28, IHH, CDH17, NGEF |
| GO:0042445 | hormone metabolic process | 2.33E-05 | 8.33E-03 | SCNN1B, BMPR1B, TTR, AFP, BCO1 |
| GO:0097746 | blood vessel diameter maintenance | 1.72E-03 | 4.48E-02 | SCNN1B, SERPINF2, CTFR |
| MM15442 | signaling by insulin receptor | 6.40E-03 | 5.23E-02 | ATP6V1B1, ATP6V1G2 |

## Discussion

FGR is currently diagnosed during the third trimester via ultrasound and fetal measurements, however, the mechanisms underlying placental insufficiency are established well before. Hence, it is imperative to the development of effective in utero treatments that we understand the intricate cellular mechanism which result in placental insufficiency at gestationally relevant time points. In the present study, we show that placental insufficiency, due to maternal nutrient restriction and increased maternal physiological stress, is associated with a reduction in transport mechanisms and vitamin synthesis including downregulation of cation/anion and amino acid transport, vitamin B6, omega3 and omega6 metabolism in the placenta. Placental metabolism of carbohydrates, glycogen synthesis, and insulin secretion is also reduced, whilst enzyme and phosphatase inhibitor activity are upregulated. Pathway enrichment analysis of differentially expressed genelists increased in the MNR placenta identified pathways that regulate DNA and RNA synthesis for both nuclear and mitochondrial translation. Similar changes in placental gene expression were observed when comparing normal placenta to the MNR placenta 5 days after *hIGF1* nanoparticle treatment, however, this is unsurprising given the short time period. When compared to sham treated MNR placentas, *hIGF1* nanoparticle treatment did increase gene expression of genes associated with transmembrane nutrient transport and decrease expression of genes associated with metabolic processes. This indicates the ability of the *hIGF1* nanoparticle to modify placental function to focus resources on fetal growth mechanisms over placental growth.

Placental insufficiency is characterized by a reduction in nutrients and oxygen reaching the fetus. Low nutrient/oxygen environments lead to increased cellular stress which in turn, lead to DNA damage, protein oxidation and lipid peroxidation [31-33]. When cells are damaged or sense stress, they activate damage repair mechanisms or induce cell death [34]. These responses alter signal transduction leading to changes in transcription factors to optimize the cells’ needs for survival or induce apoptosis if the insult is great enough. In the MNR placenta, genes that were increased were enriched for pathways related to degradation and repair of genetic information when compared to normal functioning Control placentas. These included enrichment in protein catabolism pathways, inflammatory responses, and DNA mutations at the nuclear and mitochondrial levels. Additionally, the abundance of pathways relating to repair mechanisms, particularly DNA repair mechanisms, suggest active response changes in an attempt to protect against increased cellular stress. Increased cellular stress responses in placentas collected from FGR pregnancies have previously been show in studies of human and animal pregnancies [35]. Not only do our findings recapitulate human studies showing similar changes in the delivered placentas of FGR patients [36, 37] but indicate that these changes occur throughout pregnancy in cases of placental insufficiency. Overall, these results indicate that the placenta is prioritizing cell survival over supporting fetal growth mechanisms.

With the MNR placenta experiencing increased cellular stress and prioritizing protective mechanisms for cell survival, resources for other cellular functions, such as increasing growth factors or nutrient transportation that are required to support fetal growth, become limited [35]. In the sham-treated MNR placenta, transmembrane transport pathways were downregulated when compared to normal Control placentas, likely contributing to reduced fetal weight that is characteristic of this model at the mid-pregnancy timepoint [22, 23]. Reduced placental transport capabilities also recapitulates the phenotype of the late term or delivered placenta in human FGR cases [38-40]. Downregulation of transmembrane transport mechanisms was also present in the MNR + *hIGF1* placentas when compared to Control. However, given the short time period (5 days) between treatment administration and sample collection, massive changes to nutrient transport capabilities because of treatment was not expected. In contrast, and despite the short treatment window, cell stress responses characteristic of the sham-treated MNR placentas were no longer upregulated. Instead, pathways relating to increased cell signaling and metabolism were upregulated, consistent with what is known about IGF1’s role in the placenta [41].

The ability to modify placental nutrient transport mechanisms is central to correcting aberrant fetal growth. Whilst a short, 5 day, *hIGF1* nanoparticle treatment did not result in significant changes to fetal weight [22], it did result in robust, positive changes in placental gene expression associated with nutrient supply which suggest the ability to increase fetal growth with longer treatment. When compared to sham treated MNR placentas, *hIGF1* nanoparticle treatment resulted in increased expression of genes relating to transmembrane transporter activity, particularly the ATP-binding cassette (ABC) transporter mechanism. The ABC superfamily of active transporters transport a variety of nutrients including amino acids, lipids, inorganic ions, peptides and saccharides, both across the cell membrane as well as intracellularly [42]. Placental nutrient transfer is intricately linked to placental blood flow, as the efficient exchange of nutrients and gases between the mother and fetus relies on optimal blood supply through both maternal and fetal circulations [43]. Enhanced placental blood flow ensures adequate delivery of essential nutrients and oxygen to the developing fetus, promoting healthy growth and development. We have previously shown the ability to increase fetal capillary volume density in the placenta with *hIGF1* nanoparticle treatment indicating the ability to potentially increase nutrient supply to the fetus [22]. This idea is further supported by the finding of increased expression of genes known to modulate the diameter of blood vessels and vasodilation with *hIGF1* nanoparticle treatment. Increased vasodilation would result in increased blood flow and thus increased nutrient transfer capacity.

Identifying the mechanisms of placental insufficiency during gestation allows for the identification of targets and development of effective interventions that may improve fetal development and growth, prolong pregnancies and prevent stillbirth. This study both identifies the key signaling and metabolic changes occurring in the placenta in mid-pregnancy in placenta insufficiency and increases our understanding of the pathways that intervening via increasing placental *hIGF1* expression regulates and corrects. Short-term *hIGF1* nanoparticle treatment to the placenta was able to mitigate aberrant changes in cell metabolism, nucleic acid degradation, and transmembrane nutrient transport, providing solid evidence about the ability to positively influence fetal growth trajectories with longer, more sustained treatment.

## Acknowledgments

We would like to thank Drs Craig Duvall and Mukesh Gupta for providing the co-polymer, Mrs. Kristin Lampe for assistance collecting placenta samples, and Dr Jason Puglise for assistance with RNA extractions. Additionally, we thank the University of Florida Interdisciplinary Center for Biotechnology Research Sequencing Core Facility for performing the RNA sequencing and Dr Alberto Riva for the bioinformatic analysis.

## Contributions

BND performed experiments, analyzed data and wrote manuscript. RLW conceived the study, performed experiments, analyzed data and wrote manuscript. HNJ obtained funding, conceived the study and edited manuscript.

## Ethics approval

Animal care and usage was approved by the Institutional Animal Care and Use Committee at Cincinnati Children’s Hospital and Medical Center (Protocol number 2017-0065).

## Data availability

All data needed to evaluate the conclusions in the paper are present in the paper and/or the Supplementary Materials. RNA Sequencing data has been uploaded to NCBI Geo under the accession number GSE269097.

